# Alignment-free methods for polyploid genomes: quick and reliable genetic distance estimation

**DOI:** 10.1101/2020.10.23.352963

**Authors:** Acer VanWallendael, Mariano Alvarez

## Abstract

Polyploid genomes pose several inherent challenges to population genetic analyses. While alignment-based methods are fundamentally limited in their applicability to polyploids, alignment-free methods bypass most of these limits. We investigated the use of *Mash*, a k-mer analysis tool that uses the MinHash method to reduce complexity in large genomic datasets, for basic population genetic analyses of polyploid sequences. We measured the degree to which *Mash* correctly estimated pairwise genetic distance in simulated diploid and polyploid short-read sequences with various levels of missing data. *Mash-*based estimates of genetic distance were comparable to alignment-based estimates, and were less impacted by missing data. We also used *Mash* to analyze publicly available short-read data for three polyploid and one diploid species, then compared *Mash* results to published results. For both simulated and real data, *Mash* accurately estimated pairwise genetic differences for polyploids as well as diploids as much as 476 times faster than alignment-based methods, though we found that *Mash* genetic distance estimates could be biased by per-sample read depth. *Mash* may be a particularly useful addition to the toolkit of polyploid geneticists for rapid confirmation of alignment-based results and for basic population genetics in reference-free systems with poor quality DNA.

## Introduction

By some estimates, 35% of plants are polyploid, including several essential crop species such as wheat, potato, and cotton (Wood et al. 2009). However, most population genetic methods and tools are designed for diploid markers, prompting researchers in polyploid plants to adapt novel analysis pipelines or force polyploid genomes into diploid analyses. The *in silico* reduction of polyploid genotypes to diploid ones leads to disposing of informative variation. In addition, it commonly leads to biases and systematic errors in the processing of polyploid genomes (Blischak et al. 2018). While there has been some progress in methods for the analysis of polyploid genomes (Meirmans et al. 2018; Flagel et al. 2019), many methods remain cumbersome, particularly for the analysis of high-throughput sequencing data.

One of the primary goals of population genetic analyses is to estimate population structure by determining the relatedness between individuals. In single nucleotide polymorphism (SNP) based studies, this involves three basic steps: aligning sequencing reads, calling SNPs, and estimating relatedness. Polyploid genomes present challenges to each of these steps. At the alignment step, the first issue is depth of coverage. Researchers frequently sequence genomic DNA to high depths of coverage in order to ensure full representation of the genome. However, a tetraploid genome will require twice the depth of coverage as a diploid, so will require a greater cost to achieve sufficient depth, or will result in fewer useful SNPs.

A second, related issue arises when calling SNPs, termed “allele dosage uncertainty” (Dufresne et al. 2013; Blischak et al. 2018). Next-generation sequencing platforms, particularly the commonly-used short read sequencing technologies, assemble many overlapping reads per locus, which allows the confidence to determine when a single base pair difference is a true SNP rather than a sequencing error. Diploid genomes can have a relatively high minor allele frequency (MAF) cut-off to distinguish heterozygotes from sequencing errors, but in polyploids low minor allele frequencies often represent informative variation on particular subgenomes. The random nature of sequencing depth means that it is virtually impossible to distinguish between categories of allele frequencies. For instance, an octoploid locus with alleles A and B could have a genotype ABBBBBBB, which should have a MAF of 0.125. The genotype AABBBBBB should have a MAF of 0.25. If a sequenced locus’ MAF were 0.18, for instance, it would be challenging to determine which is the true genotype. This challenge is compounded by the fact that fewer high-quality genomes have been assembled for polyploids, particularly for higher ploidy levels.

Finally, even if SNPs have been properly genotyped, calculating population-level statistics and individual relatedness offers additional challenges. Many of the common expectations of population genetic theory are built on the implicit assumption of diploidy, so many widely used statistics are not particularly useful for polyploids. For example, F-statistics are a cornerstone of population genetics. Owing to a greater number of potential heterozygous states, polyploids typically have high heterozygosity (H_s_). This results in systematically underestimating population differentiation when using F_st_ (Meirmans et al. 2018). Even where commonly used statistics may apply to polyploid datasets, there are practical concerns as well. Most bioinformatic tools require biallelic SNPs, so polyploid data analysts must produce custom analysis scripts that are challenging for reviewers to evaluate, or constrain analyses to the few tools that accommodate polyploids.

Solutions are being continuously developed to overcome the challenges of polyploid genome analysis. Alternatives to F-statistics have been developed that are less impacted by ploidy. One of the most useful is ϱ (Ronfort et al. 1998), which is equivalent to the average relatedness of individuals within populations versus the average relatedness. ϱ is comparable between ploidy levels (Meirmans et al. 2018), but has not been applied widely in empirical studies. For certain applications, unsupervised machine learning methods have shown great promise in identifying population genetic parameters using methods that are applicable to polyploids (Flagel et al. 2019). These methods are agnostic to many of the assumptions of other population genetic models, but the requirement of extensive training data mean they will not be applicable for all systems.

Currently, there are few resources that are both robust to the large degree of variation inherent to polyploid genomes, and repeatable enough to provide useful benchmarks between population genetic analyses. Since many of the challenges with polyploids come from the alignment and SNP-calling steps, an ideal solution would completely bypass these steps. Many alignment-free sequence analysis techniques have been developed in recent years, mostly using “words” or k-mers, subsamples of sequences with length k. Analyses that use k-mers add flexibility to sequence analysis and can better make use of computing power than alignment-based methods (Vinga & Almeida 2003; Leimeister et al. 2014). K-mer methods have already been applied to polyploid analyses with success (Ranallo-Benavidez et al. 2020), and so are a promising tool for population genetics in polyploid systems. There are many ways to compare k-mer decompositions of sequence data, but one of the most promising is the use of the MinHash algorithm. MinHash is a type of locality-sensitive hashing, an algorithmic technique to collapse similar input items into short strings, or hashes. MinHash then uses a Jaccard similarity coefficient to quickly assess how similar sets of hashes are. For sequence data, raw sequence reads for a sample can be decomposed into k-mer tables, which are then hashed and can be quickly compared between individuals. In practice, this process is many times faster than sequence alignment - researchers were able to cluster all of the organisms in the NCBI RefSeq database (54,118 organisms, 618 Gbp) in 35 CPU h without parallelization (Ondov et al. 2016). The dominant sequence-based MinHash implementation is known as *Mash* (Ondov et al. 2016). *Mash* and similar tools have been used widely in microbial metagenomics, but have not been incorporated into population genetics, despite their flexibility and wide applicability.

In this paper we evaluate the performance of *Mash* to estimate pairwise sample relationships within and across populations and species in polyploids. First, we simulate haploid and polyploid DNA sequences with known phylogenetic relationships, and show that *Mash* accurately recovers cladograms for these samples. Next, we remove reads from these simulated datasets to show the impact of missing data on *Mash* estimation relative to alignment-based analysts, and find that *Mash* is more robust to missing data than alignment-based estimation at estimating pairwise relationships. Finally, in several case studies, we use *Mash* to estimate kinship in reduced-representation sequencing reads for polyploid genera *Capsella* and *Reynoutria,* as well as whole-genome data in polyploid *Panicum* and diploid *Oryza*.

## Methods

### I. Simulated data

We first evaluated the performance of *Mash* using simulated haploid and polyploid sequence data. We simulated phylogenetic trees using *toytree* (Eaton 2020) in *Python 3.7* for 50 individuals using a tree height of 1e6, then simulated SNP loci following these trees by generating sequences in *ipcoal* (McKenzie & Eaton 2020). For haploid data, we simulated 1000 loci with 100bp each for 50 individuals, diverging with an effective population size of 1e5, and a recombination rate of 1e-9. For polyploid data, we simulated an allopolyploid species with two subgenomes by running two simulations based on the same tree and initial parameters, but different random seeds, then combining reads into a joint fasta file.

To simulate missing data, we used custom scripts in R to randomly remove sequence data for 5-75% of reads. We compared *Mash* and alignment-based analyses for each level of missing data.

#### Estimating genetic distance

We first calculated the true distance between individuals as the proportion of nucleotide differences (π) in the simulated sequence data. We then compared these true distances to distances inferred by alignment-based and alignment-free methods.

To provide the simplest comparison of methods, we ran a naive pipeline for both alignment-based and *Mash* methods, using the default parameters throughout. We aligned reads using *bwa* and *samtools*, then called SNPs using *bcftools* (Danecek & McCarthy 2017). We analyzed output vcfs in R using *vcfR, vegan,* and *adegenet* (Knaus & Grünwald 2017; Oskanen et al. 2007; Jombart 2008). We calculated genetic distance between samples using the *dist.genpop* function in *adegenet*, then built simple trees of these distances using *hclust* in base R, with the “complete linkage” setting.

For *Mash*, we first sketched the simulated reads using *mash sketch*, which uses minhashing to produce a reduced-complexity representation of the read set. We then calculated *Mash* distances between individuals using *mash dist*. We imported pairwise *Mash* distances into R and built trees again using *hclust.* We calculated the true distance between individuals as the proportion of nucleotide differences (π), and compared this to missing distance matrices using Mantel tests. We also compared the true tree to each tree inferred from missing data using the *comparePhylo* function in *ape* (Paradis et al. 2004).

### II. Polyploid genomes

To evaluate the performance of *Mash* on real data, we compared *Mash-*derived distance matrices to published data in population genetics studies in polyploid species from genera *Capsella* and *Reynoutria* (Cornille et al. 2016; VanWallendael et al. 2020 bioRxiv). We downloaded data for these case studies from the SRA (*Capsella*: PRJNA299253; *Reynoutria*: PRJNA574173). To account for sequencing error, we removed k-mers with counts under 2, used a k-mer length of 21, and increased the number of k-mers per sketch to 1e6. We used the same analysis tools as for simulated data, and visualized sample relationships using Principal Coordinates Analysis in R. We found that per-sample read count biased *Mash* analyses, so we randomly removed reads from the raw sequence files to standardize inputs.

While these examples are useful for ensuring that *Mash* can at least match the performance of alignment-based methods, it is not clear in these reference-free studies how well either method estimates **true** population relationships. We downloaded whole genome-sequencing data for 16 samples in *Panicum virgatum* (switchgrass; SRA Bioproject PRJNA622568) as a reference, and calculated the genetic distance between individuals using 11 million SNPs. We then simulated reduced-representation sequence data by trimming whole-genome sequences to reads that contain a typical RAD (Restriction-Associated Digest) restriction enzyme cut site (EcoRI), then dropped low-quality reads to standardize read count across samples. Using the whole-genome SNP data as “true” population relationships, we then evaluated how well *Mash* and alignment-based methods approximate these relationships. For sequence alignments, we used *bwa* (Li 2013) to align to the switchgrass v5 reference genome, then called SNPs using the Stacks *ref_map.pl* RAD-processing pipeline (Catchen et al. 2013). We called 783 SNPs from this pipeline, and estimated the genetic distance between aligned samples again using the *dist.genpop* function in *adegenet*. For *Mash* analyses, we calculated genetic distances from whole-genome sequences as well as simulated RAD sequences. We used the same parameters as for the case studies: removing k-mers with counts under 2, a k-mer length of 21, and increasing the number of k-mers per sketch to 1e6.

### III. Referenced diploid genome

To assess the performance of *Mash* on diploid data, without the implicit biases and complications of polyploid data, we also analyzed a previously published dataset of 52 rice varieties (Zheng et al. 2017). These varieties were chosen to represent the pedigrees of several elite lines, and therefore were accompanied by pedigree information for each line. This dataset allowed us to assess the performance of *Mash* in recapitulating a known relatedness structure. K-mers were sketched and analyzed as described above on a personal laptop (MacBook Pro, one 2.2Ghz 4 core Intel i7 processor).

### IV. Computing

All analyses except the referenced diploid genome were run on the high-performance computing cluster at the Institute for Cyber-Enabled Research at Michigan State University (Two 2.4Ghz 20-core Intel Xeon Gold processors). Compute time estimates were gathered by running analyses using 5 CPUs and 20GB of RAM.

## Results

### I. Simulated data

For both simulated haploid and polyploid data, we found that *Mash* correctly reconstructed phylogenetic relationships for all individuals in our N = 50 population. The genetic distances were very closely correlated to *Mash* distance (Mantel test r = 0.9606, p < 0.001) for haploid data, and only slightly less tightly correlated for polyploid samples (r = 0.9332, p < 0.001). Notably, this concordance was achieved with greatly less compute time; *Mash* analysis for these samples used 6.2 s of compute time, whereas alignment used 2893.6 s of compute time.

*Mash* analyses and alignment-based analyses differed greatly when we introduced missing data (Figures 1 & 2). Alignment-based analyses were robust to lower levels of missing data, at 5% missing data, there was a 97.2% concordance between missing and true distances. However, alignment-based distances deviated more greatly from true distances at higher levels (Figure 1). Indeed, *Mash-*based distance estimates never showed less than 50% discordance from true distances, whereas alignment-based analyses performed more poorly at levels of missing data >40%, showing a marked decrease (Figure 1.

**Figure 1:**
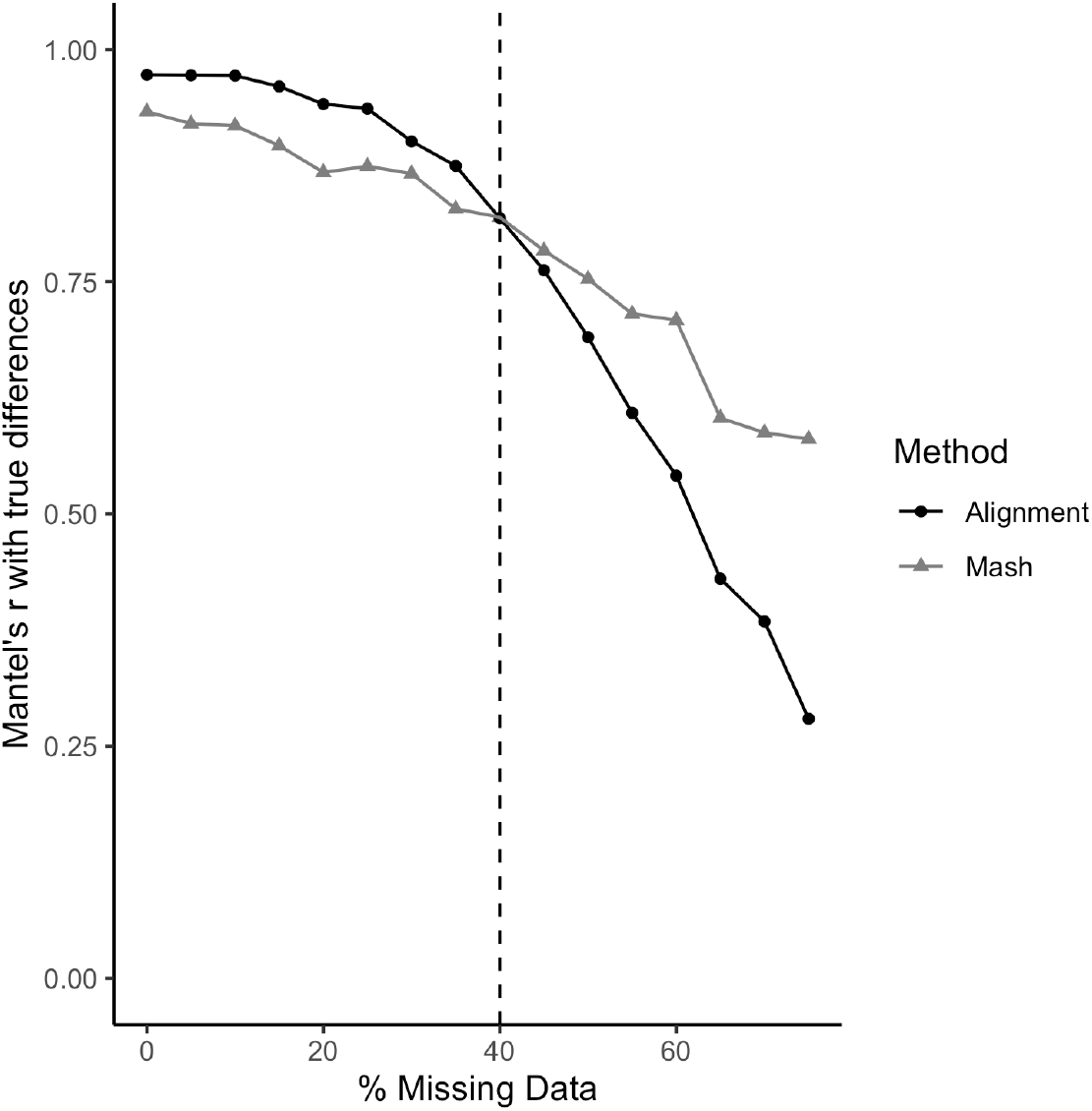
Dropoff in accuracy of pairwise genetic distance estimates for increasing levels of missing data. The black line with circles shows Mantel’s r for alignment-based methods, and the gray line with triangles shows *Mash-*based.

**Figure 2:**
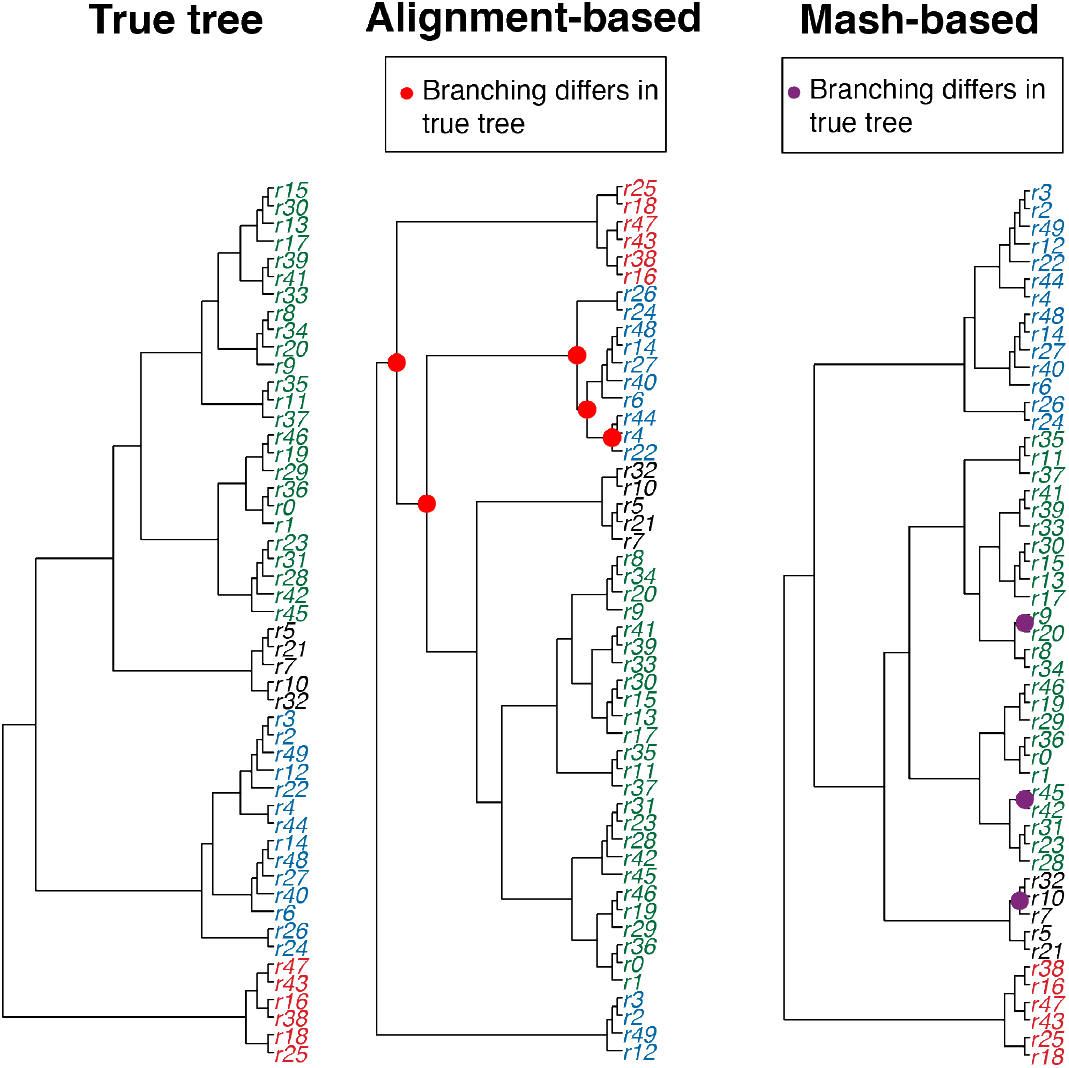
Different types of phylogenetic errors in alignment versus *Mash*-based analysis with 20% missing data of simulated polyploid data. The left panel shows the true tree used to simulate reads. Sample names are colored by major clade. The center panel shows changes in the alignment-based method, with new branches highlighted with red dots. The right panel shows changes in the *Mash* method, with new branches highlighted with purple dots.

To better understand how deviations in the pairwise genetic distance estimates can alter group assignment, we compared hierarchical clustering trees between *Mash* and alignment-based pipelines (Figure 2). While the overall Mantel’s r for *Mash* analyses at 20% missing data was slightly lower than alignment (r = 0.87 versus r = 0.94, Figure 1), the errors in genetic distance had a greater impact on phylogenetic reconstruction. The *Mash* tree showed errors in shallow nodes only, there were rearrangements even in deep nodes for the alignment-based method (Figure 2).

### II. Reduced-representation polyploid genomics

We reanalyzed data from two studies of polyploid species complexes that used genotyping-by-sequencing (Cornille et al. 2016; VanWallendael et al. 2020 bioRxiv). Using *Mash* we were able to reconstruct population groups for both studies (Figure 3 AB). As we discuss further below, there was some bias in *Mash* analyses in samples with greatly divergent read counts, but we were able to adjust by randomly trimming reads from raw sequence data. For *Capsella*, *Mash* showed five population groups, rather than the three found by ADMIXTURE, but showed high similarity to reconstructed population distributions despite using different methods (Figure 3A). We compared compute time for *Reynoutria* between *Mash* and alignment-based methods. To generate distance matrices from raw sequence reads for 25Gb of data took 0.233 compute hours in *Mash* and 108 compute hours for alignment. We could not benchmark *Capsella* alignment-based analyses, but *Mash* analyzed 44.3Gb of data in 0.410 compute hours. Since sketching is easily multithreaded, most computers can do this in a fraction of the time.

**Figure 3:**
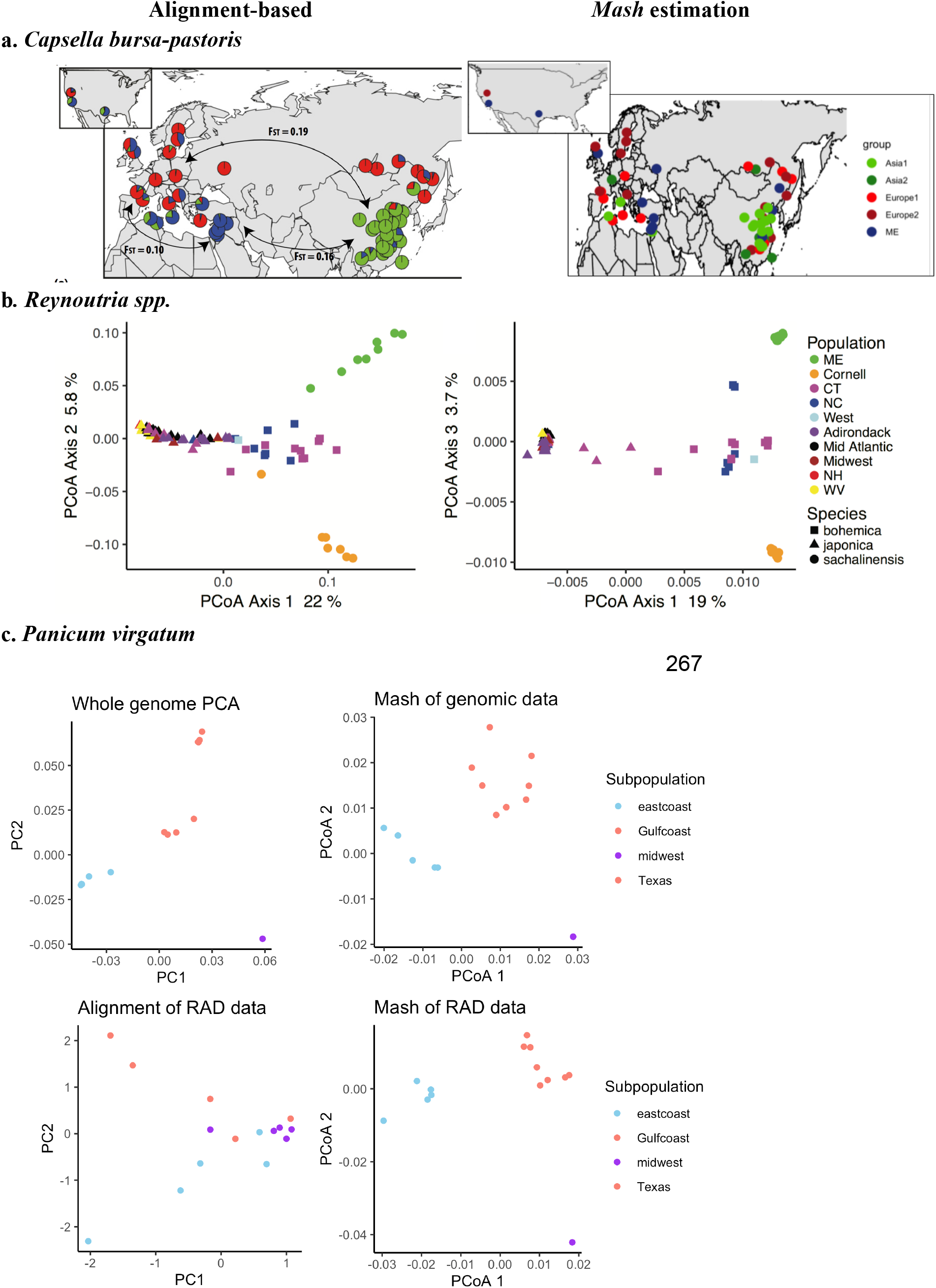
Comparisons of reference-aligned and *Mash* analyses in three genera: *Capsella, Reynoutria,* and *Panicum.* The left panels of each section show SNP-based representations of population relationships, the right panels show recreations from *Mash* analyses. a. Global diversity in *Capsella bursa-pastoris*, left panel reproduced from Cornille et al. 2016. b. Diversity in *Reynoutria spp.*, copied from VanWallendael et al. 2020. c. Whole-genome and simulated RAD data for *Panicum virgatum.* Data from NCBI SRA PRJNA622568.

**Figure 4:**
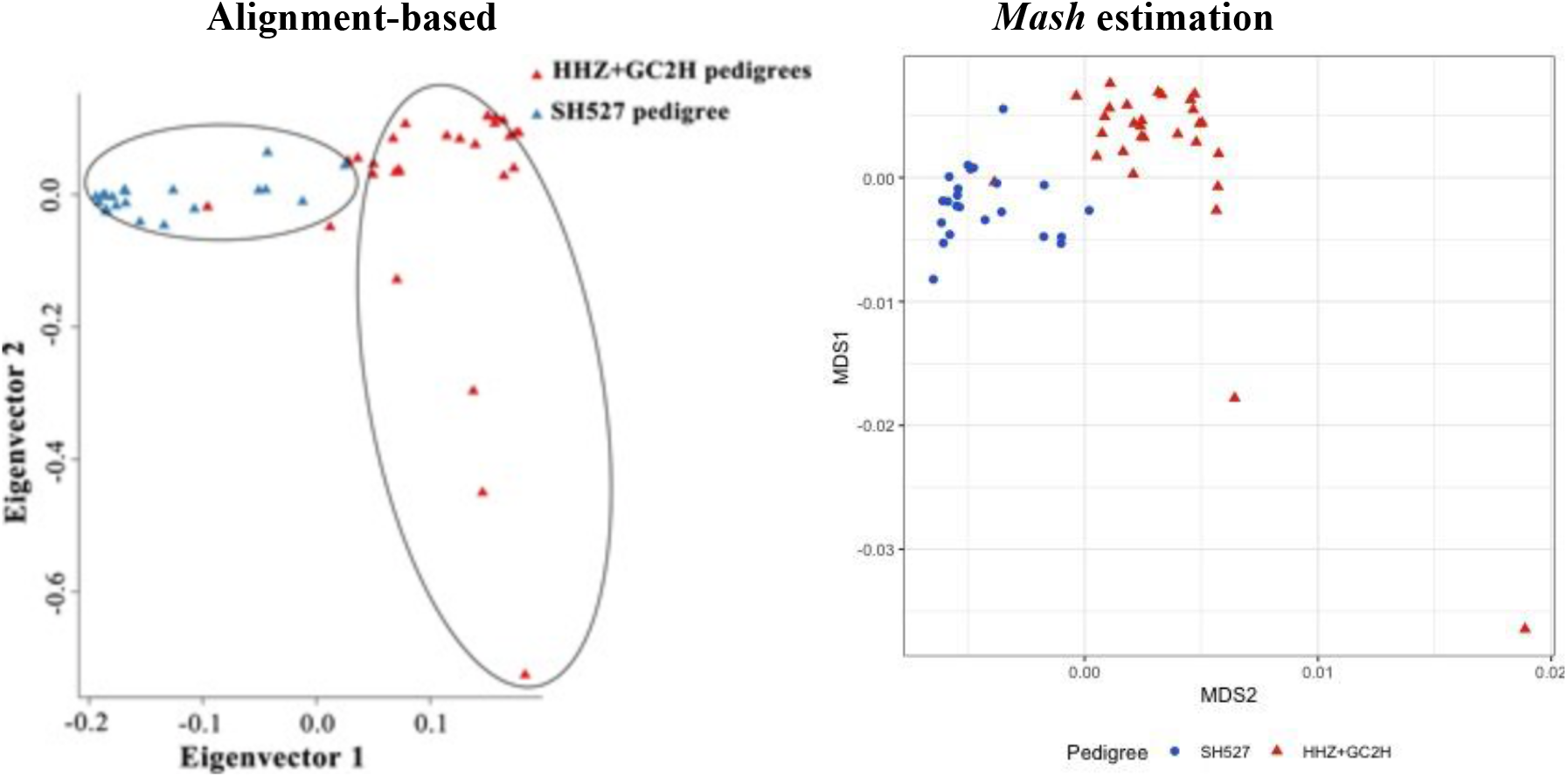
Comparison of reference-aligned and *Mash*-based scores for pedigreed O. sativa individuals. Reference-aligned plot is reproduced from Zheng et al. 2017. Note that eigenvectors are on different axes between the two plots.

To evaluate the performance of *Mash* against “known” population relationships in real sequence data, we compared the performance of *Mash* distances against alignment-based methods for simulated RAD data. We found that *Mash* effectively reproduced population relationships in *Panicum virgatum* for both whole-genome and simulated RAD data. Comparatively, the alignment-based method performed poorly, showing much less difference between known subpopulations than *Mash* of RAD sequences (Figure 3C).

### III. Whole-genome diploid analysis

When assessing diploid *O. sativa* individuals, *Mash* correctly recapitulated the known relationships between rice varieties. Patterns of population structure across the first two multidimensional scaling axes accurately separated HHZ+GC2H individuals from SH527, while maintaining outliers shown in the original plot, demonstrating the accuracy of MinHash methods in capturing patterns of variation in diploids as well as polyploids. As in previous analyses, we noted a minor effect of sequencing depth. Although this issue did not affect the ability of *Mash* to assess overall population structure, it indicates that depth correction may improve *Mash*-derived estimates of population structure. Although we did not test these methods here, other authors have explored the use of additional bias-correction steps, which improve distance calculations (*e.g.* Tang et al. 2019).

## Discussion

This study showed that *Mash* performs well at assessing genetic relationships between polyploid individuals in a fast, naive, and repeatable manner for both simulated and real data. For simulated data, we showed that *Mash* outperforms alignment-based methods in distance estimation when missing data is 40% or greater (a reasonable amount of missing data, even in whole genome sequencing projects) and is orders of magnitude faster. For real data, we have shown that *Mash* can reproduce population structure estimates in both polyploids and diploids, though per-sample read count should be standardized before analysis. *Mash* should therefore be considered a useful tool for researchers investigating population relationships in their samples, particularly when missing data is high, or when rapid population relationship assessment is necessary.

### I. Comparison with other alignment-free methods

A number of alignment-free methods using similar methods to perform k-mer hashing were reviewed by Zielezinski et al. (2019). Alternative methods are based on the length of common substrings (Ulitsky et al. 2006), micro-alignments (Leimeister et al. 2019), sequence representations based on chaos theory (Almeida et al. 2001), moments of the positions of the nucleotides (Yau et al. 2008), Fourier transformations (Yin and Yau 2015), information theory (Vinga 2014), and iterated-function systems (Almeida 2014). Of these, only *Mash* and two others were generic enough to perform protein classification, genome-based phylogenies, and detection of horizontal gene transfer (Zielezinski et al. 2019). *Mash* underperformed methods developed for closely-related organisms such as *andi* and *phylonium* at phylogenetic analysis of bacteria, but was among the top three tools for recovering tree topology in plant species (Zielezinski et al. 2019). The other two tools, *co-phylog* and *Multi-SpaM,* use some form of microalignments (Yi & Jin 2013; Dencker et al. 2020).

Several recent implementations have built on the *Mash* algorithm for specific uses. *Mashtree* is a *Mash* wrapper that automatically builds neighbor-joining trees from *Mash* outputs using QuickTree (Howe, Bateman, & Durbin, 2002; Katz et al. 2019). *Kmer-db* reports even faster compute times than *Mash*, largely through an improved k-mer hash and parallel implementation (Deorowicz et al. 2019). *sourmash* adds some functionality to *Mash* and implements scaled, rather than thresholded, numbers of retained k-mer hashes (Pierce et al. 2019). This method can enable comparisons of datasets that differ greatly in size, but slows down analysis (Pierce et al. 2019). We checked the performance of these other tools against *Mash* for the simulated data in this study, but neither *sourmash* nor *kmer-db* showed advantage over *Mash*.

### II. Potential applications for k-mer hashing based methods

As the quantity of available sequence data continues to increase exponentially, new tools that can rapidly process these data are needed (Barone et al. 2017). Locality-sensitive hashing has been used for data-intensive jobs like clustering metagenomes (Rasheed et al. 2013) and genome assembly (Berlin et al. 2015). The use of k-mer methods in molecular ecology has been comparatively limited, perhaps since historically this field has used less data-intensive approaches. However, now that genomics are more feasible for many molecular ecology researchers, k-mer hashing may solve some of the scaling issues that arise. In particular, three main directions will be facilitated by *Mash* and similar k-mer hashing methods: population genetics for polyploids, reference-free genome-wide association studies (GWAS), and genomics of ancient and otherwise degraded samples.

#### Population genetics for polyploids

Historically, population genetics for polyploids has been difficult, due to both the technical difficulty of accurately genotyping loci as well as the assumptions of diploidy in commonly used analyses and tools (Dufresne et al. 2014, Meirmans et al. 2018, Abbott et al. 2019). K-mer based methods are advantageous for this purpose, as they do not rely on accurately assessing genotype likelihood at each locus. Instead, the count data that they produce implicitly takes dosage into account, as different genotypic combinations are represented as different k-mer counts. In *Mash*, these k-mer counts are used to create distances between individuals, which can be used without further assumptions to assess differentiation in polyploid systems, particularly through eigenvector-based visualization such as principal coordinates analysis or multidimensional scaling. Further, K-mer methods represent a new path forward for assessing population differentiation, as many existing methods, including FASTSTRUCTURE (Raj et al. 2014) and ADMIXTURE (Alexander et al. 2009), assume diploidy. Future efforts should investigate the use of clustering algorithms on the distance matrices produced by *Mash* and similar methods. However, we caution the use of k-mer methods in assessing differences between mixed-ploidy populations, as they likely suffer some of the same biases as other distance-based measures (Meirmans et al. 2018).

Assessing population differentiation in polyploid systems is also complicated by the effects of polyploidy on commonly used statistics, such as Fst (Meirmans et al. 2018). Alternative measurements that are useful in polyploid systems (outlined in Meirmans et al. 2018) rely on partitioning variation within and between populations without making assumptions about the nature of individual genotypes. The multivariate analog of Fst is Φst, which relies on distance-based assessments of relatedness (Bird et al. 2011). Although we do not describe their use here, the accuracy of k-mer based methods suggests that they may also be useful for generating Φst and other genome-wide estimates between populations, particularly in polyploid systems. We suggest that future efforts be dedicated to assessing the accuracy of k-mers for this task.

To date we can find no published implementations of *Mash* or similar methods specifically for polyploid data, despite clear evidence that alignment-based methods pose significant challenges. Solutions to these challenges have mostly focused on generating higher-quality polyploid reference genomes (e.g. Edger et al. 2019) or on improving alignment-based methods (Gerard et al. 2018; Blischak et al. 2018; Clark et al. 2019). However, a recent pair of methods, *GenomeScope* and *Smudgeplot,* use a k-mer frequency-based method to estimate genome characteristics and visualize broad genome structure of polyploids (Ranallo-Benavidez et al. 2020).

#### Structural variants

Traditionally, population genetics has used a small sample of loci from a genome to infer the history of the organism. A major issue with this method has always been that genetic changes that accompany demographic changes are missed when using only relatively few neutral loci, particularly when these changes are accompanied by selection (Lowry et al. 2017). Even methods that use many millions of SNPs often miss structural changes in the genome that are increasingly linked to important evolutionary changes (Wellenreuther et al. 2019). While long-read sequencing can improve detection of structural variants, k-mers will also show this variation even in short-read data through presence and absence (Voichek & Weigel 2020). As molecular ecology studies increasingly use whole-genome coverage to assess population-level relationships, k-mer hashing can allow for vastly greater compute times and improved detection of both small and structural variation. So far, this method has mostly been applied to population genetics of microbial taxa (Ondov et al. 2016; Kachroo et al. 2019), but has the potential to be useful as more population geneticists adopt whole-genomic methods. In this study, we showed that *Mash* can accurately reconstruct population relationships from whole-genome data (Figure 3c), though we did not specifically investigate the importance of structural variants.

#### Reference-free GWAS

A breakthrough study recently showed that k-mer hashes can be used in the place of SNPs to perform reference-free GWA (Voichek & Wiegel 2020). The authors showed that k-mer based GWAS outperformed SNP-based methods, particularly in confidence of detection of causal variants and in the number of phenotypes caused by structural variants (Voichek & Wiegel 2020). Reference-free GWA is not completely new, but early attempts used computationally inefficient methods that are not scalable to large, complex genomes (Lees et al. 2016). As k-mer counts accurately represent complex polyploid genotypes in an efficient way, they pave the way for large-scale GWA analyses in polyploid systems. For example, just as k-mer counts replace genotypes in the linear mixed models commonly used for GWAS, k-mer-derived distance matrices can replace the relatedness matrix that is used to control for population structure in the same analyses. Critically, neither of these methods rely on an assumption of diploidy.

#### Genomics of ancient and degraded samples

A major finding in this study is that *Mash* greatly outperforms alignment-based methods when samples have missing data. This leads to similar issues as those posed in polyploid genomes. Since most phylogenetic analyses use Bayesian methods, not distance-based hierarchical clustering, this may not be an issue in that field. Phylogenetic analyses have indeed been found to be largely robust to missing data (Hovmöllar et al. 2013), however, estimation of specific population genetic parameters can be more impacted (Chattopadhyay et al. 2014). Ancient DNA studies are particularly prone to problems with missing data, since long-degraded samples often cannot yield full SNP coverage. Recent aDNA studies have included samples with as much as 40% (Marcus et al. 2020) or 50% missing data (Anava et al. 2020), while others removed SNPs with missingness >10% (Sikora et al. 2014), leading to a greatly reduced genomic coverage. Since these studies are particularly challenging to replicate given the barriers to sample collection, it is especially concerning that missing data may interfere with conclusions. K-mer hashing analysis may offer a confirmatory method for aDNA studies.

## Conclusions

*Mash* and similar alignment-free methods hold great promise for overcoming some of the challenges in both polyploid and diploid genomics. In situations where alignments are particularly fraught, such as in systems with a high level of missing data, large genomic arrangements, or data-heavy whole-genome comparisons, the simplicity and speed of these methods may offer at minimum a confirmation of alignment-based results. Other methods will need to be tested against *Mash* to determine which applies for particular population genetic studies. Similarly, combining machine learning with k-mer based analyses may hold great promise for fast and reliable population genetic estimation.

## Acknowledgements

Much thanks to Rafael Gutaker for the initial idea to use *Mash* for polyploid genomics. We would like to also thank the authors of Cornille et al. 2016 and Zheng et al. 2017 for use of their sequences from the SRA, as well as HudsonAlpha for *Panicum* sequences.

## Data Accessibility

Data and code for all analyses will be made available on Github: https://github.com/avanwallendael/mash_sim. External datasets were downloaded from the NCBI SRA Bioprojects PRJNA622568, PRJNA299253, and PRJNA574173.

